# Fides: Reliable Trust-Region Optimization for Parameter Estimation of Ordinary Differential Equation Models

**DOI:** 10.1101/2021.05.20.445065

**Authors:** Fabian Fröhlich, Peter K. Sorger

## Abstract

Ordinary differential equation (ODE) models are widely used to describe biochemical processes, since they effectively represent mass action kinetics. Optimization-based calibration of ODE models on experimental data can be challenging, even for low-dimensional problems. However, reliable model calibration is a prerequisite for uncertainty analysis, model comparison, and biological interpretation. Multiple hypotheses have been advanced to explain why optimization based calibration of biochemical models is challenging, but there are few comprehensive studies that test these hypotheses, likely because tools for performing such studies are also lacking.

We implemented an established trust-region method as a modular Python framework (`fides`) to enable systematic comparison of different approaches to ODE model calibration involving various Hessian approximation schemes. We evaluated `fides` on a set of benchmark problems for which real experimental data are available. Unexpectedly, we observed high variability in optimizer performance among different implementations of the same algorithm. Overall, `fides` performed most reliably and efficiently. Our investigation of possible sources of poor optimizer performance identified drawbacks in the widely used Gauss-Newton, BFGS and SR1 Hessian approximations. We address these drawbacks by proposing a novel hybrid Hessian approximation scheme that enhances optimizer performance and outperforms existing hybrid approaches. We expect `fides` to be broadly useful for ODE constrained optimization problems and to enable future methods development.

**Availability:** `fides` is published under the permissive BSD-3-Clause license with source code publicly available at https://github.com/fides-dev/fides. Citeable releases are archived on Zenodo. Code to reproduce results presented in this manuscript is available at https://github.com/fides-dev/fides-benchmark.

## 1 Introduction

Mass action biochemical systems can be accurately described in the continuous approximation (i.e., with a large number of molecules per reaction compartment) by ordinary differential equation (ODE) models. Although cells are not well-mixed systems, ODE modeling can be highly effective in describing biochemical processes in both eukaryotic and prokaryotic cells [1]. To ensure these ODEs recapitulate and predict experimental observations, model parameters must be estimated from data. This estimation problem can be formulated as optimization problem, where the objective function describes the discrepancy between a given solution to the ODE and experimental data. Minimizing this discrepancy can be computationally demanding due to the numerical integration required when evaluating the objective function and its derivatives [2]. Optimized parameter values are often used to initialize model analysis such as uncertainty quantification via the profile likelihood or sampling approaches [3, 4]. Similarly, parameter optimization is required when models are compared based on goodness of fit, using measures such as AIC or BIC, or when other complexity penalizing methods are applied [5, 6]. Thus, reliably and efficiently finding robust solutions to the optimization problem is of critical importance in many aspects of ODE model analysis.

In general, the optimization problem for ODE models is non-convex, resulting in few theoretical guarantees of convergence when numerical optimization is employed. It is therefore typically necessary to rely on empirical evidence to select appropriate optimization algorithms for any specific class of problems [2, 7]. The optimization problem for ODE models of biochemical systems belongs to an uncommon class of problems having four characteristic properties: (i) the optimization problem is often ill-posed due to parameter non-identifiability; (ii) with tens to hundreds of estimated parameters, they are computationally intensive, but do not qualify as high-dimensional problems in the optimization literature as, e.g., problems of that size do not raise concerns about storage of the Hessian in memory; (iii) computation time of numerically solving the optimization problem is dominated by evaluation of the objective function and its derivatives, i.e., the computation time for the proposed parameter update itself is negligible; (iv) since models are inexact and experimental data is noisy, the residual values between simulation and data may be much larger than zero, even at the global minimum of the optimization problem (such problems are commonly called non-zero residual problems). The existing benchmarks for general purpose optimization, such as the CUTE(r/st) [8–10] set of benchmarks, do not cover models having these four characteristics. Thus, domain-specific benchmarks need to be considered.

For a broad set of biochemical ODE models, trust-region methods initialized from a large number of random initial parameter values have performed well [11, 12]. Trust-region methods use local (quadratic) approximations of the objective function to propose parameter updates and then iteratively refine the local neighborhood in which the local approximation is expected to adequately recapitulate the shape of the true objective function, i.e. the trust-region [13]. Popular implementations of trust-region methods are available in the MATLAB optimization toolbox and the scipy Python optimization module. However, for a large fraction of problems, including low dimensional biochemical models with as few as 20 parameters, these optimizers do not consistently converge to parameter values that yield similar values for the objective function [11], strongly suggesting that the global optimum - and possibly even a local optimum - has not been reached. For example, the benchmark study by Hass *et al.* performed optimization for a model based on work of Fujita *et al.* [14], which describes Endothelial Growth factor mediated activation of the Protein Kinase B pathway (commonly known as AKT pathway). They found that the difference in negative log-likelihood between the best and second best parameter values was > 20, exceeding the statistical threshold for model rejection according to AIC and BIC criteria [15]. For such problems, model selection is a challenge, because poor optimizer performance may erroneously lead to the rejection of a model.

More generally, the presence of “optimal” solutions of parameter values that yield inconsistent values for the objective function (Fig 1A) can indicate either (i) that optimization converged on a few of the many critical points (local minima, saddle points) (Fig 1B right) or (ii) that optimization terminated before convergence to any (local) minimum was achieved (Fig 1B left). However, many of the problems of interest in biochemistry and cell biology have multiple local minima, which can be a result of curvature of the model manifold [16]. In these cases, repeated convergence of multiple optimization runs on a small set of similar objective function values may not represent a problem with the optimization problem itself, but rather arise from model non-identifiability. This setting contrasts with the situation in which the objective function values are inconsistent, despite a large number of runs (setting rigorous thresholds for what can be considered “consistent” is a tricky problem in and of itself, which we revisit later in this manuscript). In such a situation, it is unclear whether optimization is non-convergent or the objective function is very “rugged” [2] with many local minima, not all of which have been identified.

**Fig 1:**
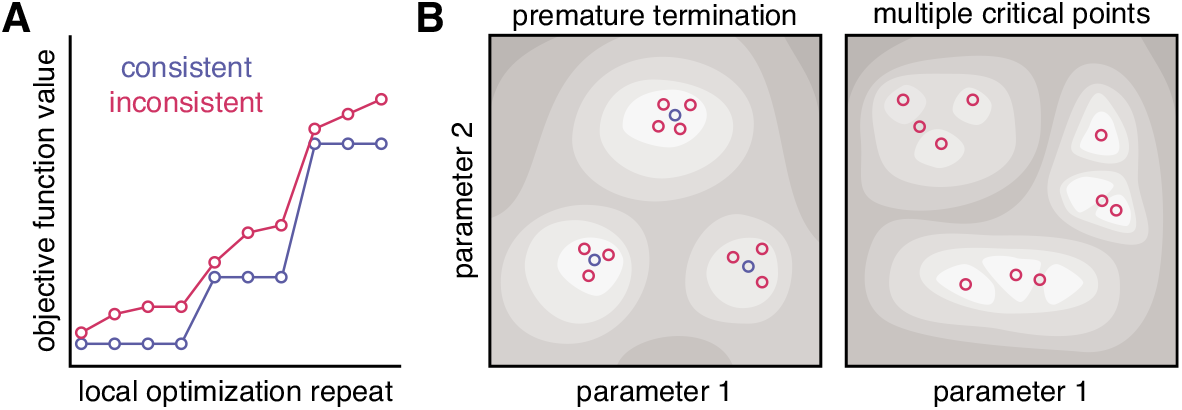
Illustration of final objective function values consistency and possible objective function landscapes. **A:** Waterfall plot with examples of consistent (blue) and inconsistent (red) final objective function values. **B:** Possible objective function landscapes that could explain the waterfall plots in A.

Non-convergent optimization results can be the consequence of noisy model simulations, in which lax integration tolerances result in inaccurate numerical evaluation of the objective function value and its gradient [17]. Inaccurate gradients often result in poor parameter update proposals, slowing the search in parameter space. Inaccurate objective function values may result in incorrect rejection of parameter updates, both of which can erroneously suggest convergence to a minimum and thus lead to premature termination of optimization. For example, Tönsing *et al.* [17] found that optimization runs with similar, but inconsistent objective function values were often located in the neighborhood of the same local minima, consistent with the idea that optimizer runs had been prematurely terminated. They suggested that a nudged elastic band method, which aims to identify shortest connecting paths between optima, might be effective in improving consistency. Premature termination can also arise from problems with the optimization method itself. Dauphin *et al.* [18] found that saddle points are prevalent in the objective functions of neural network models and that optimization methods that do not account for directions of negative curvature may perform poorly in the vicinity of saddle points. However, neither the prevalence of saddle points nor their impact on premature optimizer termination have been investigated in the case of biochemical ODE models. Lastly, Transtrum *et al.* [16] have suggested that the use of Gauss-Newton Hessian approximations may not work well for sloppy models. Sloppiness is encountered when the objective function Hessian has a broad eigenvalue spectrum, suggesting non-identifiability of parameters and an ill-posed optimization problem. Sloppiness is believed to be a universal property of biochemical models [19]. However, the geodesic acceleration proposed by Transtrum *et al.* [16] to address the limitations of Gauss-Newton Hessian approximation has not been widely adopted, likely due to the implementation complexity and computational cost of determining directional second-order derivatives.

Overall, the results described above show that early optimizer termination is a recurrent issue and may have a variety of causes. Yet, a comprehensive evaluation of this issue on many benchmark problems, as well as development and testing of methods to identify or resolve the underlying causes are missing. In principle, this could be addressed by adapting optimization algorithms. For example, it might be possible to resolve issues with the Gauss-Newton Hessian approximation by using alternative approximation schemes, such as the Broyden-Fletcher-Goldfarb-Shanno (BFGS) [20–23] scheme. Issues with saddle points could be resolved by employing symmetric rank-one (SR1) [24] approximations that account for negative curvature directions. However, many optimization algorithms were written decades ago, are difficult to customize or extend, and do not provide the user with statistics about individual optimization traces. These limitations make it difficult to diagnose problems with optimization and to resolve them with algorithmic improvements.

To tackle these and other challenges associated with optimization of biochemical models based on ODEs, this paper re-implements a standard trust-region algorithm in Python and uses it to investigate several hypotheses about causes and potential solutions for poor optimizer performance. We find that the use of an inaccurate Hessian approximation is one important contributor to poor optimization performance and propose a novel hybrid Hessian approximation scheme. We demonstrate that this scheme outperforms existing approaches on a set of benchmark problems.

## 2 Materials and Methods

For the purpose of this study, we considered four different optimizers that all implement the interior-trust-region algorithm proposed by Coleman and Li [25]: `fmincon`, referring to the MATLAB function of the same name and with `trust-region-reflective` as algorithm and `ldl-factorize` as subproblem algorithm, `lsqnonlin`, referring to the MATLAB function of the same name, `ls_trf`, referring to the scipy function `least_squares` with `trf` algorithm, and `fides`, the new implementation developed in the current manuscript. Below we describe algorithmic and implementation details of `fides` (a summary is provided in Table 1), and of the benchmark problems we used to evaluate these algorithms.

**Table 1:** Feature overview for different trust-region optimization implementations. The non least-squares column indicates whether the method is applicable to non least-squares problems. The free column indicates whether the implementation is freely available or proprietary software.

| Optimizer | Subspace | Non least-squares | BFGS/SR1 | Programming Language | free |
| --- | --- | --- | --- | --- | --- |
| lsqnonlin | $\mathcal{S}_{2D}$ | □ | □ | MATLAB | □ |
| fmincon | $\mathcal{S}_{2D}$ | ✓ | □ | MATLAB | □ |
| ls_trf | $\mathbb{R}^{n_\theta}, \mathcal{S}_{2D}$ | □ | □ | Python | ✓ |
| fides | $\mathbb{R}^{n_\theta}, \mathcal{S}_{2D}$ | ✓ | ✓ | Python | ✓ |

### 2.1 Model Formulation

When applied to a biochemical system, an ODE model describes the temporal evolution of abundances of *n*_*x*_ different molecular species *x*_*i*_. The temporal evolution of x is determined by the vector field *f* and the initial condition x_0_:

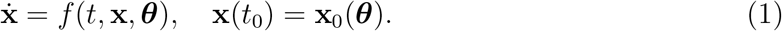

Both *f* and x_0_ depend on the unknown parameters 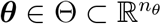 such as catalytic rates or binding affinities. Restricting optimization to the parameter domain Θ can constrain the parameter search space to values that are realistic based on physicochemical theory and helps prevent numerical integration failures associated with extreme parameter values. For most problems, Θ is the tensor product of scalar search intervals (*l*_*i*_, *u*_*i*_) with lower and upper bounds *l*_*i*_ < *u*_*i*_ that satisfy *l*_*i*_, *u*_*i*_ ∈ ℝ ∪ {− ∞, ∞} for every parameter *θ*_*i*_.

Experiments usually provide information about observables y which depend on abundances x and parameters ***θ***. A direct measurement of x is usually not possible. Thus, the dependence of observables on abundances and parameters is described by

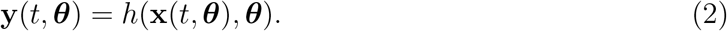

#### Implementation in this Study

All methods described here use CVODES from the SUNDIALS suite [26] for numerical integration of model equations. CVODES is a multi-step implicit solver for stiff- and non-stiff ODE initial value problems.

### 2.2 Optimization Problem

To generate models useful in the study of actual biological systems, model parameters ***θ*** must be inferred from experimental data, which are typically incomplete and subject to measurement noise. A common assumption is that the measurement noise for *n*_*t*_ time-points *t*_*j*_ and *n*_*y*_ observables *y*_*i*_ is additive, independent and normally distributed for all time-points:

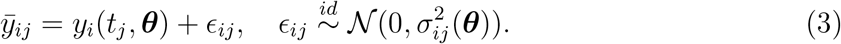

Thus, model parameters can be inferred from experimental data by maximizing the likelihood, yielding a maximum likelihood estimate (MLE). However, the evaluation of the likelihood function involves the computation of several products of large terms, which can be numerically unstable. Thus, the negative log-likelihood

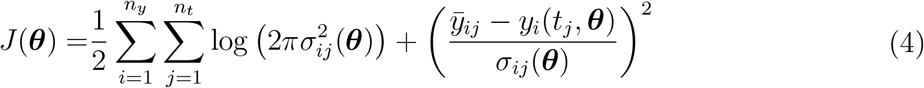

is typically used as objective function that is minimized. As the logarithm is a strictly monotonously increasing function, the minimization of *J* (***θ***) is equivalent to the maximization of the likelihood. Therefore, the corresponding minimization problem

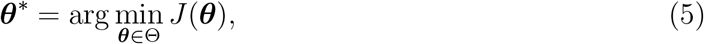

will infer the MLE parameters. If the noise variance 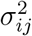 does not depend on the parameters ***θ***, the objective function (4) has a weighted least-squares formulation. As we discuss later, properties of a least-squares formulation can be exploited in specialized optimization methods. Optimizers that do not require least-squares structure can also work with other noise models [27].

#### Implementation in this Study

For the MATLAB optimizers `fmincon` and `lsqnonlin`, the objective function and its derivatives were evaluated using data2dynamics [28] (commit `b1e6acd`), which was also used in the study by Hass *et al.* [11]. For the Python optimizers `ls_trf` and `fides`, the objective function and its derivates were evaluated using AMICI [29] (version 0.11.23) and pyPESTO (version 0.2.10).

### 2.3 Trust-Region Optimization

Trust-region methods minimize the objective function *J* by iteratively updating parameter values ***θ***_*k*+1_ = ***θ***_*k*_ + ∆***θ***_*k*_ according to the local minimum

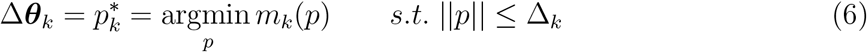

of an approximation *m*_*k*_ to the objective function. ∆_*k*_ is the trust-region radius that restricts the norm of parameter updates. The optimization problem (6) is known as the trust-region subproblem. In most applications, a local, quadratic approximation

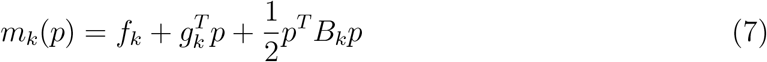

is used, where *f*_*k*_ = *J* (***θ***_*k*_) is the value, *g*_*k*_ = ▽*J* (***θ***_*k*_) is the gradient and *B*_*k*_ = ▽^2^*J* (***θ***_*k*_) is the Hessian of the objective function evaluated at ***θ***_*k*_.

The trust-region radius ∆_*k*_ is updated in every iteration depending on the ratio *ρ*_*k*_ between the predicted decrease 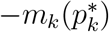 and actual decrease in objective function value 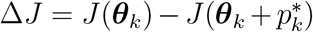 [13]. The step is accepted if *ρ*_*k*_ exceeds some threshold *μ* ≥ 0. When boundary constraints (on parameter values) are applied, the predicted decrease is augmented by an additional term that accounts for the parameter transformation (see Section 2.6) [25].

#### Implementation in this Study

All optimizers evaluated in this study use *μ* = 0 as acceptance threshold. They all increase the trust-region radius ∆_*k*_ by a factor of 2 if the predicted change in objective function value is accurate (*ρ*_*k*_ > 0.75) and the local minimum is at the edge of the trust region (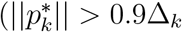 for `fmincon`, `lsqnonlin` and `fides`, 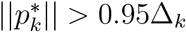 for `ls_trf`). All optimizers decrease the trust-region radius if the predicted change in objective function value is inaccurate (*ρ* < 0.25), but `fmincon`, lsqnonlin and `fides` set the trust-region radius to 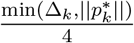, while `ls_trf` sets it to 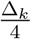.

When the predicted objective function decrease is negative 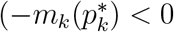, i.e., an increase in value is predicted) `fides` and `ls_trf` set *ρ*_*k*_ to 0.0. Positive values for 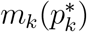 may arise from the augmentation accounting for boundary constraints. Setting *ρ*_*k*_ to 0.0 prevents inadvertent increases to ∆_*k*_ or step acceptance when 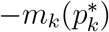 and ∆*J* are both negative. However, in contrast to `fides`, `ls_trf` does not automatically reject respective step proposals and only does so if ∆*J* < 0. `fmincon` and lsqnonlin only reject step proposals with ∆*J* < 0, but do not take the sign of 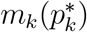 into account when updating ∆_*k*_.

When the objective function cannot be evaluated – for ODE models this is typically the result of an integration failure – all optimizers decrease the trust-region radius by setting it to 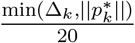 (`fmincon` and `lsqnonlin`), 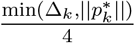 (`fides`) or 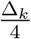 (`ls_trf`). These subtly nuanced differences in the implementation are likely the result of incomplete specification of the algorithm in the original publication [25]. In particular, handling of 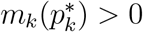, which does not occur for standard trust-region methods, was not described and developers needed to independently work out custom solutions.

### 2.4 Hessian Approximation

Constructing the local approximation (7) that defines the trust-region subproblem (6) requires the evaluation of the gradient *g*_*k*_ and Hessian *B*_*k*_ of the objective function at the current parameter values ***θ**_k_*. While the gradient *g*_*k*_ can be efficiently and accurately computed using first order forward or adjoint sensitivity analysis [30], it is computationally more demanding to compute the Hessian *B*_*k*_ [31]. Therefore several approximation schemes have been proposed that approximate *B*_*k*_ using first order sensitivity analysis. In the following we will provide a brief description of approximation schemes considered in this study, an overview of schemes and their characteristics is provided in Table 2.

**Table 2:** Overview of properties of different Hessian approximation schemes. BFGS is the Broyden-Fletcher-Goldfarb-Shannon algorithm. SR1 is the Symmetric Rank-one update. GN is the Gauss-Newton approximation. SSM is the Structured Secant Method. TSSM is the Totally Structured Secant Method. FX is the hybrid method proposed by Fletcher and Xu [39]. GNSBFGS is the Gauss-Newton Structured BFGS method. The *construction column* indicates whether pointwise evaluation is possible or whether iterative construction is necessary. The *positive semi-definite* column indicates whether the approximation preserves positive semi-definiteness given a positive semi-definite initialization.

| Scheme | Construction | Positive Semi-Definite | Convergence Requirement | Requires Least-Squares |
| --- | --- | --- | --- | --- |
| **BFGS** | iterative | ✓ | ✓ | □ |
| **SR1** | iterative | □ | ✓ | □ |
| **GN** | pointwise | ✓ | $\ \mathbf{r}(\boldsymbol{\theta}^*)\ = 0$ | ✓ |
| **SSM** | pointwise + iterative | □ | ✓ | ✓ |
| **TSSM** | pointwise + iterative | □ | ✓ | ✓ |
| **FX** | pointwise + iterative | ✓ | ✓ | ✓ |
| **GNSBFGS** | pointwise + iterative | ✓ | $\lambda_{\min}(\nabla^2 J(\boldsymbol{\theta}^*)) > 0$ | ✓ |

#### Gauss-Newton Approximation

The Gauss-Newton (GN) approximation 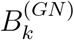 is based on a linearization of residuals *r*_*ij*_

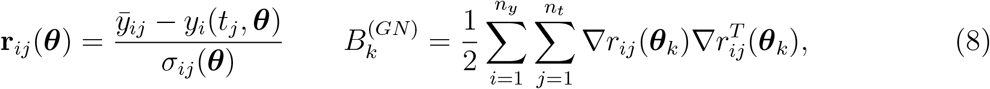

which yield a symmetric and positive semi-definite approximation to *B*_*k*_ and does not account for negative curvature. At the maximum likelihood estimate, *B*^(*GN*)^ is equal to the negative empirical Fisher Information Matrix assuming *σ*_*ij*_ does not depend on parameter values ***θ***.

For parameters dependent *σ*, the log(*σ*) term in (4) cannot be assumed to be constant, which results in a non least-squares optimization problem. For non least-squares problems, the adequateness and formulation of the GN approximation is not well established. Thus, Raue [32] proposed to transform the problem into a least-squares form by introducing additional error residuals 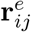 and adding a corresponding correction to the Gauss-Newton approximation *B*^(*GN*)^ from (8), yielding *B*^(*GNe*)^:

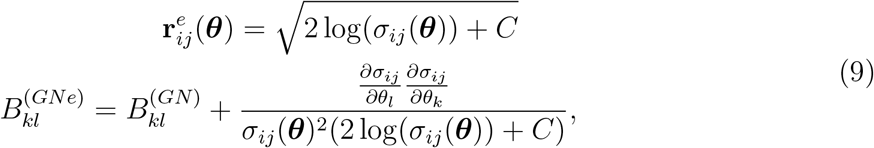

where *C* is some arbitrary, but sufficiently large constant so that 2 log(*σ*_*ij*_ (***θ***)) + *C* > 0. This condition ensures that residuals are real-valued and the approximation *B*^(*GNe*)^ is positive semi-definite, but inherently makes optimization a non-zero residual problem. Adding the constant *C* to residuals adds a constant to the objective function value and, thus, neither influences its gradient and Hessian nor the location of its minima. However, *C* does enter the GNe approximation, with unclear implications. Instead, Stapor *et al.* [31] suggested that one ignores the second order derivative of the log(*σ*) term in (4), which corresponds to the limit lim_*C*→∞_ *B*^(*GNe*)^ = *B*^(*GN*)^.

#### Iterative Approximations

In contrast to the GN approximation, Broyden-Fletcher-Goldfarb-Shanno (BFGS) or Symmetric Rank-one (SR1) are iterative approximation schemes, in which the approximation in the next step

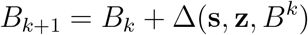

is constructed based on the approximation in the current step *B*_*k*_ and some update ∆(*x, s, B^k^*), where s = ∆***θ***_*k*_ and **z** = *g*_*k*+1_ − *g*_*k*_.

The BFGS update scheme

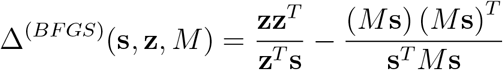

guarantees a positive semi-definite approximation as long as a curvature condition **z**^***T***^ s > 0 is satisfied and the initial approximation 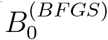 is positive semi-definite [13]. Thus, the update scheme is usually only applied with line search methods that guarantee satisfaction of the curvature condition by selecting the step length according to (strong) Wolfe conditions [13]. However, BFGS can also be used in trust-region methods by rejecting updates when the curvature condition is not satisfied, although this invalidates some theoretical convergence guarantees [13].

The SR1 update scheme

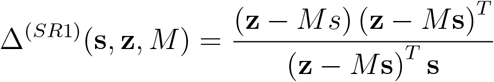

can also yield indefinite approximations, incorporating negative curvature information, and has no step requirements beyond ensuring that the denominator of the update is non-zero.

#### Structured Secant Approximations

The accuracy of the GN approximation depends on the magnitude of the residuals, since the approximation error is the sum of products of the residuals and the residual Hessians

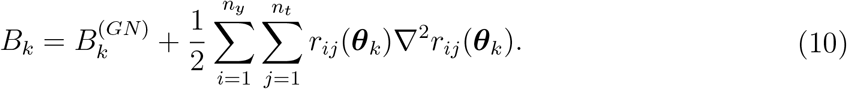

For non-zero residual problems, in which there are indices *i, j* such that *r*_*ij*_(***θ***^*^) ≫ 0, or in the presence of strong non-linearities, the second order term in (10) does not vanish and the GN approach is known to perform poorly; it can even diverge [33, 34]. This issue is addressed in structured secant methods [33, 35, 36], which combine the pointwise GN approximation with an iterative BFGS approximation *A*_*k*_ of the second order term. In the Structured Secant Method (SSM) [36], the matrix *A*_*k*_ is update using a BFGS scheme:

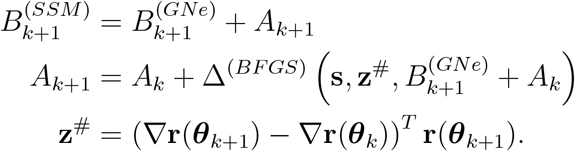

Similarly, the Totally Structured Secant Method (TSSM) [37] scales *A*_*k*_ with the residual norm, to mimic the product structure in the second order term in (10), and, accordingly, scales the update to *A*_*k*_ with the inverse of the residual norm:

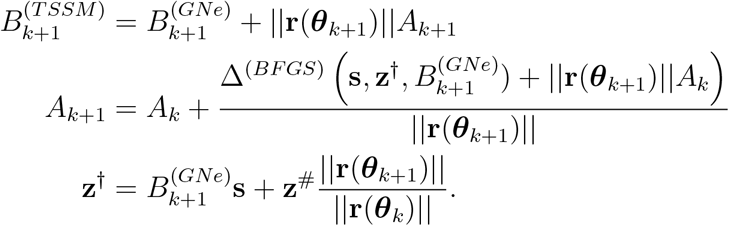

Despite the use of a BFGS updating scheme, the SSM and TSSM approximations do not preserve positive semi-definiteness, as the matrix 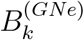 is updated at every iteration without any additional safeguards. Structured secant approximations have been popularized by the NL2SOL toolbox [38], but are not featured in the standard optimization libraries in MATLAB or Python.

#### Hybrid Schemes

Other hybrid schemes can dynamically switch between GN approximations and iterative updates when some metric indicates that the considered problem has non-zero residual structure. For example, Fletcher and Xu [39] proposed an approach to detect non-zero residuals by computing the normalized change in the residual norm and applying BFGS updates when the change is smaller than some tolerance *ϵ*_*FX*_:

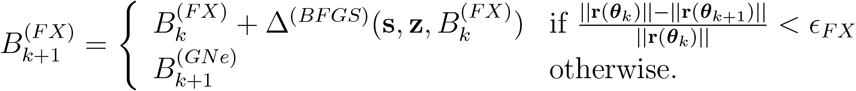

As 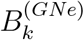 is positive semi-definite and the BFGS updates preserve this property, the FX approximations are always positive semi-definite.

To address the issue of possibly indefinite approximations in the SSM and TSSM approaches, Zhou and Chen proposed a Gauss-Newton structured BFGS method (GNSBFGS) [34].

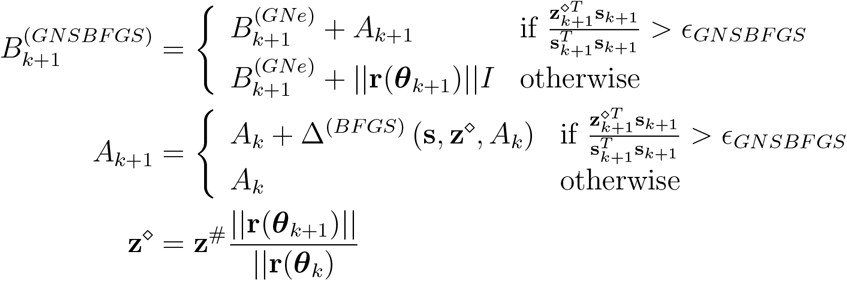

that combines the TSSM approach with the dynamic updating of the FX approach. As sum of two positive semi-definite matrices, 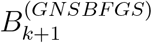 is always positive semi-definite. The authors demonstrate that the term 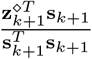 plays a similar role as the 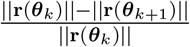 term in the FX algorithm. However, they only prove convergence if the tolerance 2*ϵ*_*GNSBFGS*_ is smaller than the smallest eigenvalue *λ*_min_(∇^2^*J* (***θ***^*^)) of the Hessian at the optimal parameters and if ∇^2^*J* (***θ***^*^) is positive definite. This condition is not met for problems with singular Hessians, which are often observed for non-identifiable problems.

#### Implementation in this Study

`fmincon` and `lsqnonlin` were only evaluated using the GNe approximation, as implemented in data2dynamics. `ls_trf` can only be applied using the GNe approximation. `Fides` was evaluated using BFGS and SR1 using respective native implementations in addition to GN and GNe, as implemented in AMICI. We used the default value of *C* = 50 for the computation of GNe in both data2dynamics and AMICI. We provide implementations for BFGS, SR1, SSM, TSSM, FX and GNSBFGS schemes in `fides`. FX, SSM, TSSM and GNSBFGS were applied using GNe, as they require a least-squares problem structure. Hyperparameters *ϵ*_*FX*_ = 0.2 and *ϵ*_*GNSBFGS*_ = 10^−6^ were picked based on recommended values in respective original publications.

### 2.5 Solving the Trust-Region Subproblem

In principle, the trust-region subproblem (6) can be solved exactly [13]. Moré proposed an approach using eigenvalue decomposition of *B*_*k*_ [40]. Yet, Byrd *et al.* [41] noted the high computational cost of this approach and suggested an approximate solution by solving the trust-region problem over a two dimensional subspace 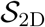, spanned by gradient *g*_*k*_ and Newton 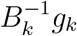 search directions, instead of 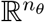. Yet, for objective functions requiring numerical integration of ODE models, the cost of eigenvalue decomposition is generally negligible for problems involving fewer than 10^3^ parameters.

A crucial issue for the two-dimensional subspace approach are problems with indefinite (approximate) Hessians. For an indefinite *B*_*k*_, the Newton search direction may not represent a direction of descent. This can be addressed by dampening *B*_*k*_ [13], but for boundary-constrained problems additional considerations arise and require the identification of a direction of strong negative curvature [25].

#### Implementation in this Study

`fmincon` and `lsqnonlin` implement optimization only over 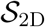, where the Newton search direction is computed using preconditioned direct factorization. For preconditioning and direct factorization, `fmincon` and `lsqnonlin` employ Cholesky and QR decomposition respectively, which both implement dampening for numerically singular *B*_*k*_. `fides` and `ls_trf` implement optimization over 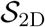 (denoted by 2D in text and figures) and 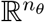 (denoted by `ND` in text and figures). For `ls_trf`, we specified tr solver=”lsmr” for optimization over 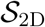. To compute the Newton search direction, `ls_trf` and `fides` both use least-squares solvers, which is equivalent to using the Moore-Penrose pseudoinverse. `fides` uses the direct solver `scipy.linalg.lstsq`, with `gelsd` as LAPACK driver, while `ls_trf` uses the iterative, regularized `scipy.sparse.linalg.lsmr` solver. In `fides`, the negative curvature direction of indefinite Hessians is computed using the eigenvector to the largest negative eigenvalue (computed using `scipy.linalg.eig`).

### 2.6 Handling of Boundary Constraints

The trust-region region subproblem (6) does not account for boundary constraints, which means that *θ*_*k*_ + ∆*θ*_*k*_ may not satisfy these constraints. For this reason, Coleman and Li [25] introduced a rescaling of the optimization variables depending on how close the current values are to the parameter boundary *∂*Θ. This rescaling is realized through a vector

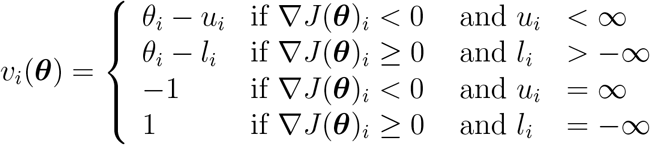

which yields transformed optimization variables

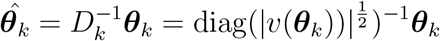

and a transformed Hessian

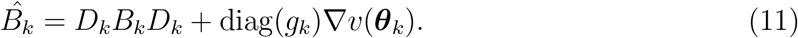

As the second term in (11) is positive semi-definite, this handling of boundary constraints can also regularize the trust-region sub-problem, although this is not its primary intent.

Coleman and Li [25] also propose a stepback strategy in which solutions to (6) are reflected at the parameter boundary *∂*Θ. Since this reflection defines a one-dimensional search space, the local minimum can be computed analytically at negligible computational cost. To ensure convergence, ∆*θ*_*k*_ is then selected based on the lowest *m*_*k*_(*p*) value among (i) the reflection of *p*\* at the parameter boundary (ii) the constrained Cauchy step, which is the minimizer of (7) along the gradient that is truncated at the paramtere boundary and (iii) *p*\* truncated at the parameter boundary.

#### Implementation in this Study

`fmincon`, `lsqnonlin` and `ls_trf` all implement rescaling, but only allow for a single reflection at the boundary [42]. In contrast, `fides` implements rescaling and also allows for a single or arbitrarily many reflections until the first local minimum along the reflected path is encountered.

### 2.7 Optimizer Performance Evaluation

To evaluate optimization performance, Hass *et al.* [11] computed the success count *γ*, which represents the number of “successful” optimization runs that reached a final objective function value sufficiently close (difference smaller than some threshold *τ*) to the lowest objective function value found by all methods, and divided that by the time to complete all optimization runs *t*_*total*_, a performance metric that was originally introduced by Villaverde *et al.* [43]. In this study, we replaced *t*_*total*_ with the total number of gradient evaluations across all optimization runs *n*_*grad*_ for any specific optimization setting. The resulting performance metric *ϕ* = *γ/n*_*grad*_ ignores differences in computation time for gradients having different parameter values and prevents computer or simulator performance from influencing results. This provides a fairer evaluation of the algorithm or method itself and is particularly relevant when optimization is performed on computing clusters with heterogeneous nodes or when different number of threads are used to parallelize objective function evaluation, as it was the case in this study. Since *φ* ignores potentially higher computation times for step-size computation, we confirmed that step-size computation times were negligible as compared to numerical integration of model and sensitivity equations. For all trust-region optimizers we studied, the number of gradient evaluations was equal to the number of iterations, with the exception of `ls_trf` which only uses objective function evaluations when a proposed step is rejected. Thus, 1/*n*_*grad*_ is equal to the average number of iterations for the optimization to converge (divided by the number of optimization runs, which is 10^3^ in all settings). We therefore refer to *ν* = 1/*n*_*grad*_ as the convergence rate. Performance *ϕ* is equal to the product of *γ* and *ν*.

To calculate *γ*, we used a threshold of *τ* = 4, which corresponds to the upper limit of the objective function value in cases in which a model cannot be rejected according to the BIC [44]. Similar to Hass *et al.* [11], we found that changing this convergence threshold did not have a significant impact on performance comparison, but provide analysis for values *τ* = 0.05 (threshold used by Hass *et al.*, divided by two to account for difference in objective function scaling, Fig S1) and *τ* = 20 (the threshold for rejection according to AIC and BIC [15], Fig S2) in the Supplementary Material.

### 2.8 Extension of Boundary Constraints

For some performance evaluations, we extended parameter boundaries. Even though initial points are usually uniformly sampled in Θ, we did not modify the locations of initial points when extending bounds. The *Schwen* problem required a different approach in which the bounds for the parameter `fragments` were not modified, as values outside the standard bounds were implausible.

As previously reported [11], extending boundaries can expose additional minima having globally lower objective function values. Thus, success count *γ* for optimization settings with normal boundaries were computed using the lowest objective function *J*_min_ found among all settings excluding those with extended boundaries. *γ* for optimization settings using extended boundaries were computed using the minimum of *J*_min_ and the lowest objective function value found for that particular setting.

### 2.9 Statistical Analysis of Optimizer Traces

During the statistical analysis of optimizer traces, we quantified several numerical values derived from numerical approximation of matrix eigenvalues with limited accuracy (due, for example, to limitations in floating point precision). It was therefore necessary to account for this limitation in numerical accuracy:

#### Singular Hessians

To numerically assess matrix singularity of Hessian approximations, we checked whether the condition number, computed using the numpy function `numpy.linalg.cond`, was larger than the inverse of the floating point precision *ϵ* =1/numpy.spacing(1).

#### Negative eigenvalues

To numerically assess whether a matrix has negative eigenvalues we computed the smallest (*λ*_min_) and largest (*λ*_max_) eigenvalues of the untransformed Hessian approximation *B*_*k*_ using `numpy.linalg.eigvals` and checked whether the smallest eigenvalue had a negative value that exceeded numerical noise *λ*_min_(*B*_*k*_) < −*ϵ* · |*λ*_max_(*B*_*k*_)|.

### 2.10 Implementation

`fides` is implemented as modular, object-oriented Python code. The subproblem, subproblem solvers, stepback strategies and Hessian approximations all have class-based implementations, making it easy to extend the code. Internally, `fides` uses SciPy [45] and NumPy [46] libraries to store vectors and matrices and perform linear algebra operations. To ensure access to state-of-the-art simulation and sensitivity analysis methods, we implemented an interface to `fides` in the parameter estimation toolbox pyPESTO, which uses AMICI [29] to perform simulation and sensitivity analysis via CVODES [26]. This approach also enabled import of biological parameter estimation problems specified in the PEtab [47] format.

### 2.11 Benchmark Problems

To evaluate the performance of different optimizers, we considered 13 out of the 20 different benchmark problems (Table 3) introduced by Hass *et al.* [11]; these problems were reformulated in the PEtab [47] format. The selection of benchmark problems was based on whether or not all of the functionality required for the PEtab import was supported and validated in AMICI and pyPESTO and whether the problems could be encoded in SBML and PEtab formats. For example the *Hass* problem was excluded since it includes negative initial simulation time, which is not supported by PEtab. Other models were excluded for the following reasons: *Raia* because it involves state-dependent value for *σ*, which is unsupported by AMICI; *Merkle* and *Sobotta* due to missing SBML implementations; *Swameye* because it includes spline functions which are not supported by SBML; *Becker* because it involves multiple models, which is not supported by pyPESTO; and *Chen* because forward sensitivity analysis is prohibitively computationally expensive for this model. All benchmarks problems were previously published and included experimental data for model calibration as described by Table 3, which also provides a brief summary of numerical features of the various benchmarks.

**Table 3:** Summary of problem characteristics for benchmark examples, as characterized by Hass *et al.* [11].

| Problem | $n_\theta$ | $n_x$ | $n_y \cdot n_t$ | sloppy | identifiable |
| --- | --- | --- | --- | --- | --- |
| *Bachmann* | 113 | 36 | 541 | ✓ | □ |
| *Beer* | 72 | 4 | 27132 | ✓ | □ |
| *Boehm* | 9 | 8 | 48 | ✓ | ✓ |
| *Brannmark* | 22 | 9 | 43 | ✓ | □ |
| *Bruno* | 13 | 7 | 77 | □ | ✓ |
| *Crauste* | 12 | 5 | 21 | ✓ | □ |
| *Fiedler* | 19 | 6 | 72 | ✓ | □ |
| *Fujita* | 19 | 9 | 144 | ✓ | □ |
| *Isensee* | 46 | 25 | 687 | ✓ | □ |
| *Lucarelli* | 84 | 43 | 1755 | ✓ | □ |
| *Schwen* | 30 | 11 | 286 | ✓ | □ |
| *Weber* | 36 | 7 | 135 | ✓ | □ |
| *Zheng* | 46 | 15 | 60 | ✓ | □ |

A more detailed description of the biochemical systems described by these models is available in the supplemental material of the study by Hass *et al.* [11].

### 2.12 Simulation and Optimization Settings

We encountered difficulties reproducing some of the results described by Hass *et al.* [11] and therefore repeated evaluations using the latest version of data2dynamics. We deactivated Bessel correction [11] and increased the function evaluation limit to match the iteration limit. Relative and absolute integration tolerances were set to 10^−8^. The maximum number of iterations for optimization was set to 10^5^. Convergence criteria were limited to step sizes with a tolerance of 10^−6^, and `ls_trf` code was modified such that the convergence criteria matched the implementation in other optimizers. For all problems, we performed 10^3^ optimizer runs. To initialize optimization, we used the initial parameter values provided by Hass *et al.* [11].

### 2.13 Parallelization and Cluster Infrastructure

Optimization was performed on the O2 Linux High Performance Compute Cluster at Harvard Medical School running SLURM. Optimization for each model and each optimizer setting was run as a separate job. For MATLAB optimizers, optimization was performed using a single core per job. For Python optimizers, execution was parallelized on up to 12 cores. We observed severe load balancing issues due to skewed computational cost across optimization runs for *Bachmann*, *Isensee*, *Lucarelli* and *Beer* models. To mitigate these issues, optimization for these models was parallelized over 3 threads using pyPESTOs `MultiThreadEngine` and simulation was parallelized over 4 threads using openMP multithreading in AMICI, resulting in a total parallelization over 12 threads. For the remaining models, optimization was parallelized using 10 threads without parallelization of simulations. Wall-time for each job was capped at 30 days (about 1 CPU year), which was only exceeded by the GNSBFGS and FX Hessian approximations for the *Lucarelli* problem after 487 and 458 optimization runs respectively and also by the SSM Hessian approximation for the *Isensee* problem after 973 optimization runs. Subsequent analysis was performed using partial results for those settings.

## 3 Results

### 3.1 Validation and Optimizer Comparison

The implementation of trust-region optimization involves complex mathematical operations that can result in error-prone implementations. To validate the trust-region methods implemented in `fides`, we compared the performance of optimization using GN and GNe schemes against implementations of the same algorithm in MATLAB (`fmincon`, `lsqnonlin`) and Python (`ls_trf`). In the following, the employed subspace solvers and Hessian approximation methods are denoted by *implementation subspace*/*hessian*.

We found that `fides 2D/GN` (blue) and `fides 2D/GNe` (orange) were the only methods that had non-zero performance (*ϕ* > 0) for all 13 benchmark problems (Fig 2A), with small performance differences between the two methods (0.72 to 1.12 fold difference, average 0.96). This established `fides 2D/GN` as good reference implementation. We will, therefore, report the performance of other methods relative to `fides 2D/GN` (Fig 2B). The `ls_trf` method outperformed `fides 2D/GN` on three problems (1.54 to 22.4-fold change; *Boehm*, *Crauste*, *Zheng*; purple arrows), had similar performance on one problem (1.15-fold change; *Fiedler*), exhibited worse performance on four problems (0.02 to 0.55-fold change; *Brannmark*, *Bruno*, *Lucarelli*, *Weber*) and did not result in successful runs (zero performance *ϕ* = 0) for the remaining five problems (*Bachmann*, *Beer*, *Fujita*, *Isensee*, *Schwen*). Decomposing performance improvements *ϕ* into increases in convergence rate *ν* (Fig 2C) and success count *γ* (Fig 2D) revealed that increase in *ϕ* was primarily due to higher *ν*, which was observed for all but four problems (*Bachmann*, *Bruno*, *Isensee*, *Schwen*). However, in most cases, improvements in *ν* were canceled out by larger decreases in *γ*.

**Fig 2:**
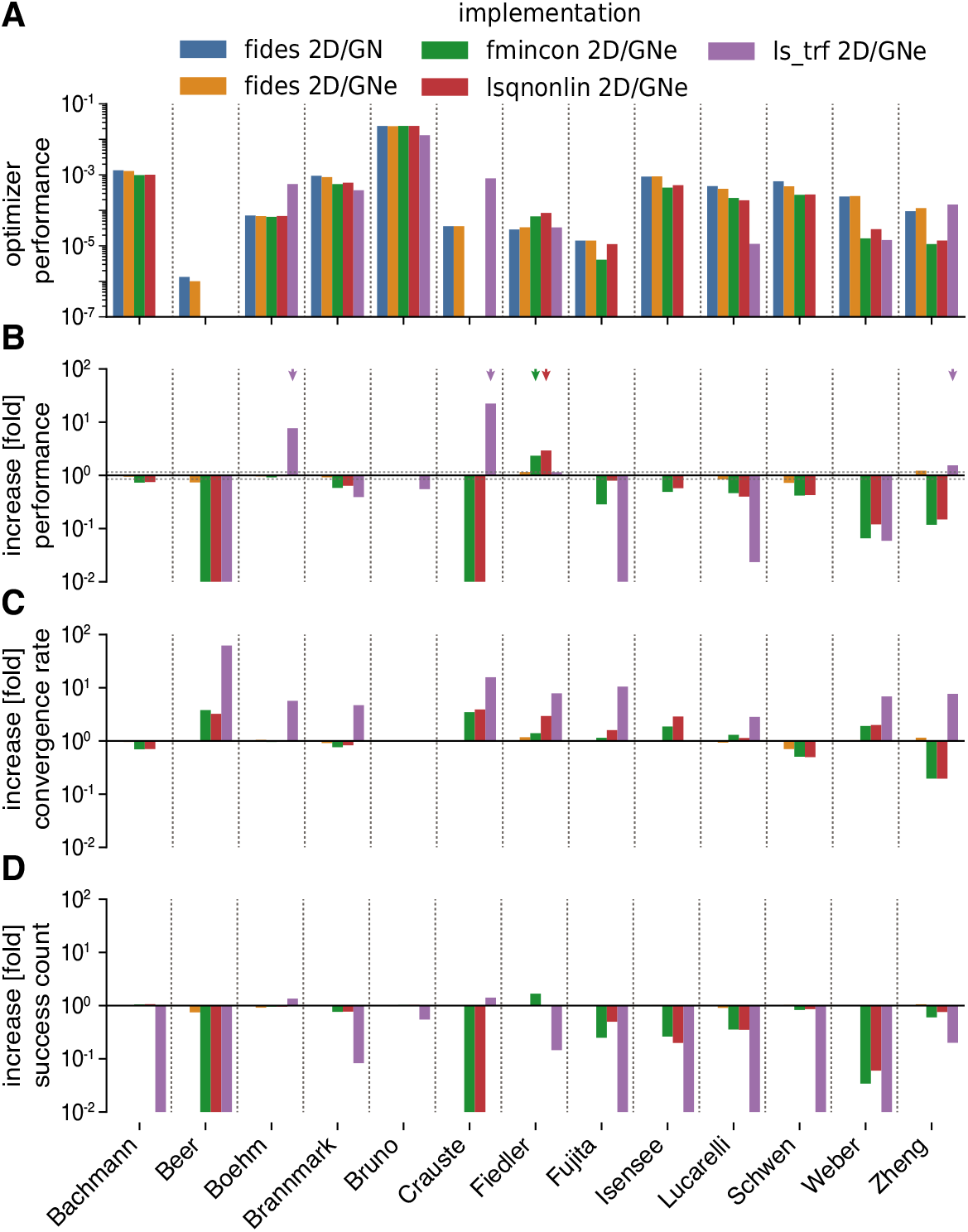
Comparison of MATLAB and Python optimizers. Colors indicate optimizer setting and are the same in all panels. **A:** Performance comparison (*ϕ*), absolute values. **B:** Performance comparison (*ϕ*), values relative to `fides 2D/GN`. **C:** Increase in convergence count *γ*, values relative to `fides 2D/GN`. **D:** Increase in convergence rate *ν*, values relative to `fides 2D/GN`.

For `fmincon` (green) and `lsqnonlin` (red), *ϕ* was higher for one problem (2.34 to 2.94 -fold change; *Fiedler*; red/green arrow), similar for two problems (0.92 to 1.00 fold change, *Boehm*, *Bruno*) and worse for the remaining 10 problems (0.00 to 0.80 fold change, *Bachmann*, *Brannmark*, *Fujita*, *Isensee*, *Lucarelli Schwen*, *Weber*, *Zheng*), with zero performance on two problems (*Beer*, *Crauste*) (Fig 2B). Since we observed similar *ϕ* for `fides 2D/GN` (blue) and `fides 2D/GNe` (orange) on all problems, the differences in *ϕ* between `fides 2D/GN` and the other implementations are unlikely to reflect use of GNe as opposed to a GN scheme. Instead we surmised that the differences were due to discrepancies in implementation of the Newton direction (this would explain the similarity for the two identifiable problems (*Boehm*, *Bruno*, Table 3)).

Overall, these findings demonstrated that trust-region optimization implemented in `fides` was more than competitive with the MATLAB optimizers `fmincon` and `lsqnonlin` and the Python optimizer `ls_trf`, outperforming them on a majority of problems. Simultaneously, our results demonstrate a surprisingly high variability in optimizer performance among methods that implement the same algorithm. This variability may explain some of the conflicting findings in previous studies that assumed distinct implementations would have similar performance [48, 49].

### 3.2 Parameter Boundaries and Stepback Strategies

One of the few changes in implementation that we deliberately introduced into the `fides` code was to allow multiple reflections during stepback from parameter boundary conditions [25]. In contrast, `ls_trf`, `lsqnonlin` and `fmincon` only allow a single reflection [42]. The modular design and advanced logging capabilities of `fides` make it straightforward to evaluate the impact of such modifications on optimizer performance *ϕ* and arrive at possible explanations for observed differences. For example, when we evaluated `fides 2D/GN` with single (orange) and multi-reflection (dark-green, Fig 3A-C) implementations and correlated changes to *ϕ* with statistics of optimization trajectories (Fig 3D,E), we found that the single reflection performance *ϕ* was reduced on four problems (0.59 to 0.65-fold change; *Beer*, *Lucarelli*, *Schwen*, *Zheng*; orange arrows Fig 3A). Lower performance was primarily due to a decrease in convergence rate *ν* (Fig 3B,C). We attributed this behavior to the fact that a restriction on the number of reflections lowered the predicted decrease in objective function values for reflected steps. This, in turn, increased the fraction of iterations in which stepback yielded constrained Cauchy steps (Pearson’s correlation coefficient *r* = −0.85, p-value *p* = 2.3 · 10^−4^, Fig 3D) as well as the average fraction of boundary-constrained iterations (*r* = −0.82, p=6 · 10^−4^, Fig 3E), both slowing convergence.

**Fig 3:**
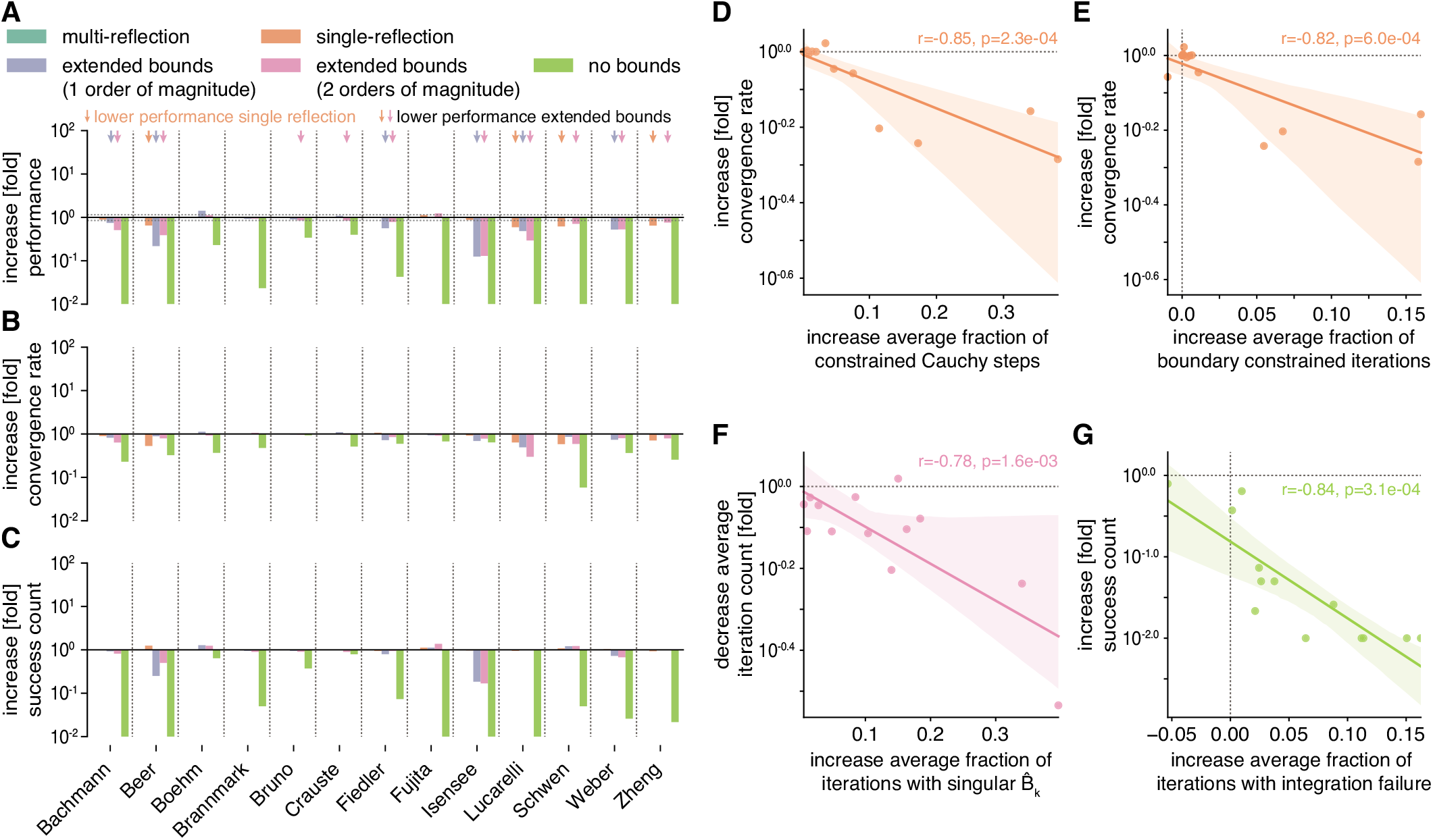
Evaluation of stepback strategies. Colors indicate optimizer setting and are the same in all panels. All increases/decreases are relative to multi-reflection `fides 2D/GN` with normal bounds. **A:** Performance comparison (*ϕ*). **B:** Increase in convergence count *γ*. **C:** Increase in convergence rate *ν*. **D:** Association between increase in average fraction of constrained Cauchy steps with respect to total number of boundary constrained iterations and increase in *ν* for the single-reflection method. **E:** Association between increase in average fraction of boundary constrained iterations and increase in *ν* for the single-reflection method. **F:** Association between increase in average fraction of iterations with a numerically singular transformed Hessian 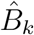 and decrease in *γ* for the multi-reflection method for `fides 2D/GN` with bounds extended by two orders of magnitude. **G:** Association between increase in average fraction of iterations with integration failures and increase in *γ* for multi-reflection `fides 2D/GN` without bounds.

A naive approach to addressing issues with parameter boundaries is to extend or remove the parameter bounds. Thus, we repeated optimization with `fides 2D/GN` (multi-reflection) with parameter boundaries extended by one (blue) or two (pink) orders of magnitude or completely removed (light green). We found that extending boundaries by one order of magnitude reduced *ϕ* for 6 problems (0.12 to 0.74 fold change; *Bachmann*, *Beer*, *Fiedler*, *Isensee*, *Lucarelli*, *Weber*; blue arrows Fig 3A) and extending boundaries by two orders of magnitude reduced *ϕ* for an additional 4 problems (0.13 to 0.85 fold change; *Bruno*, *Crauste*, *Schwen*, *Zheng*; pink arrows Fig 3A). We found that decreased *ϕ* was primarily the result of lower *ν* (Fig 3B), which we attributed to a larger fraction of iterations in which the transformed Hessian 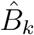 was singular (*r* = −0.78, *p* = 1.6 · 10^−3^, Fig 3F). Removing boundaries decreased *ϕ* for all problems, a result of lower values of *γ*, which we attributed to higher fraction of iterations with integration failures (*r* = −0.82, *p* = 5.7 · 10^−4^, Fig 3G).

These findings demonstrate the importance and difficulty of choosing appropriate optimization boundaries, since excessively wide boundaries may lead to frequent integration failures and/or the creation of an ill-conditioned trust-region subproblem. In contrast, too narrow boundaries may not include the global optimum. When managing boundary constraints the use of multi-reflection as compared to single-reflection yields a small performance increase, albeit significantly smaller than the variation we observed (in the previous section) between different implementations of the same optimization algorithm.

### 3.3 Iterative Schemes and Negative Curvature

To further study the positive effect of improving the conditioning of trust-region subproblems on the optimizer performance *ϕ*, we carried out optimization using BFGS and SR1 Hessian approximations. BFGS and SR1 can yield full-rank Hessian approximations, resulting in well-conditioned trust-region subproblems, even for non-identifiable problems. Moreover, in contrast to GN, these approximations converge to the true Hessian under mild assumptions [13]. The SR1 approximations can also account for directions of negative curvature and might therefore be expected to perform better when saddle-points are present.

We compared *ϕ* for `fides 2D/GN` (blue), `fides 2D/BFGS` (orange) and `fides 2D/SR1` (green) (Fig 4A) and found that `fides 2D/BFGS` failed to reach the best objective function value for one problem (*Beer*) and `Fides 2D/SR1` for two problems (*Beer*, *Fujita*). Compared to `fides 2D/GN`, *ϕ* for `fides 2D/BFGS` was higher on three problems (2.02 to 9.88 fold change; *Boehm*, *Fiedler*, *Schwen*; orange arrows) and four problems for `fides 2D/SR1` (1.32 to 7.12 fold change; *Boehm*, *Crauste*, *Fiedler*; green arrows), it was lower for a majority of the remaining problems (BFGS 7 of 13 problems, 0.05 to 0.49 fold change; SR1 8 of 13 problems, 0.07 to 0.78 fold change). Decomposing *ϕ* into improvements in convergence rate *ν* (Fig 4B) and improvements in success counts *γ* (Fig 4C) revealed that SR1 improved *ν* for 7 problems (2.23 to 5.97 fold change; *Beer*, *Boehm*, *Crauste*, *Fiedler*, *Fujita*, *Weber*, *Zheng*; green arrows Fig 4B). Out of these 7 problems, BFGS improved *ν* for only four problems (2.33 to 6.63 fold change; *Beer*, *Boehm*, *Fiedler*, *Zheng*; orange arrows Fig 4B). We found that for both approximations, the increase in *ν* was correlated with the change in average fraction of iterations without trustregion radius (∆_*k*_) updates (BFGS: *r* = −0.8, *p* = 1.1 · 10^−3^; SR1: *r* = −0.61, *p* = 2.8 · 10^−2^, Fig 4D). ∆_*k*_ is not updated when the predicted objective function decrease is in moderate agreement with the actual objective function decrease (0.25 < *ρ*_*k*_ < 0.75, see Section 2.3), likely a result of inaccurate approximations to the Hessian. Thus, higher convergence rate *ν* of SR1 and BFGS schemes was likely due to more precise approximation of the objective function Hessian.

**Fig 4:**
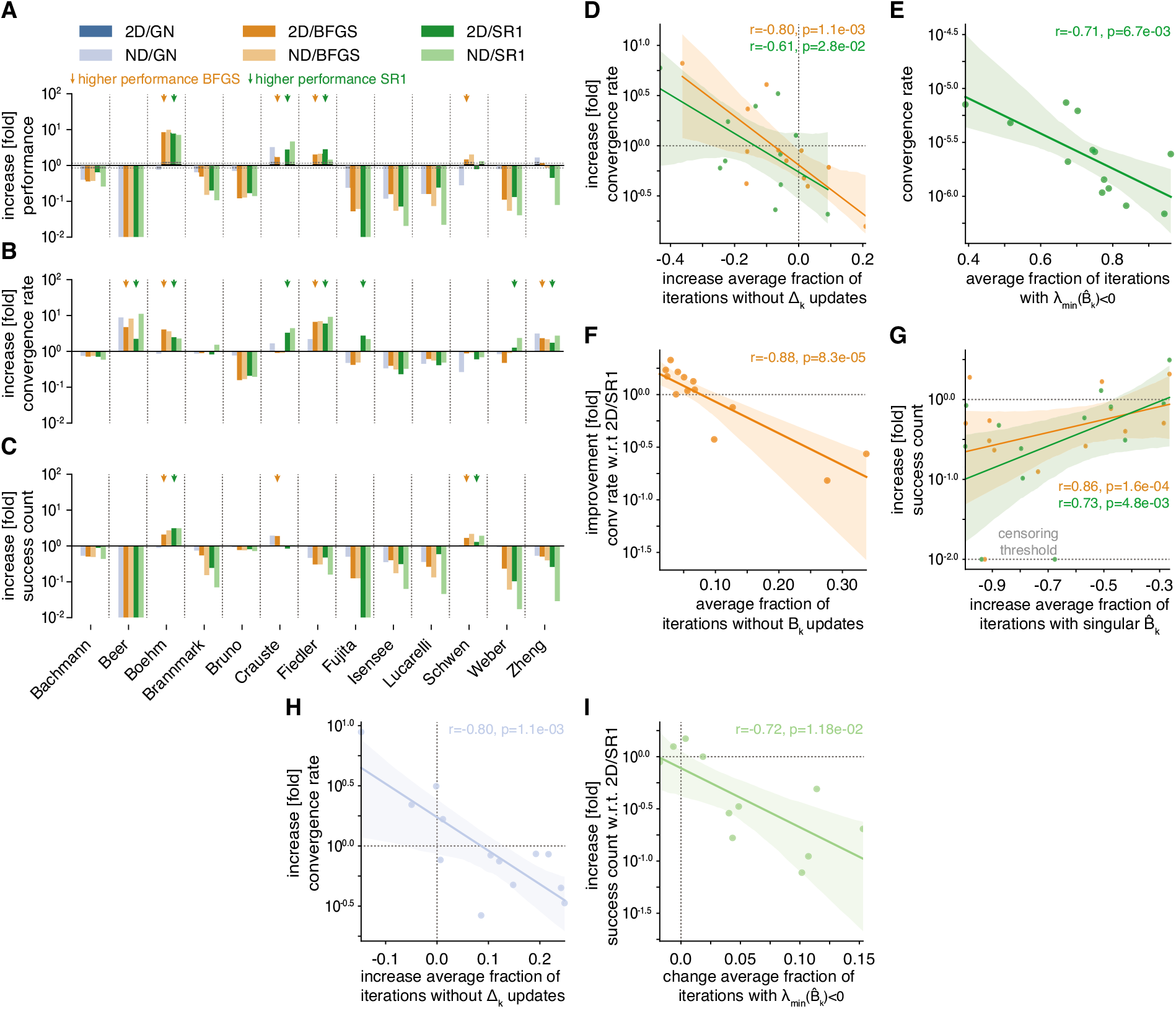
Evaluation of iterative Hessian approximation schemes. Color scheme is the same in all panels. All increases/decreases are relative to `fides 2D/GN` unless otherwise noted. **A:** Performance comparison (*ϕ*). **B:** Increase in convergence count *γ*. **C:** Increase in convergence rate *ν*. **D:** Association between increase in average fraction of iterations without updates to the trust-region radius ∆_*k*_ and increase in *ν* for `fides 2D/SR1` (green) and `fides 2D/BFGS` (orange). **E:** Association between average fraction of iterations where smallest eigenvalue *λ*_min_ of the transformed Hessian 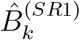 is negative and *ν* for `fides 2D/SR1`. **F:** Association between fraction of iterations without updates to *B*_*k*_ and increase in *ν* relative to `fides 2D/SR1` for `fides 2D/BFGS`. **G:** Association between fraction of iterations with a numerically singular transformed Hessian 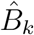 and increase in *γ* for `fides 2D/SR1` (green) and `fides 2D/ BFGS` (orange). Change in *γ* was was censored at a threshold of 10^−2^ to visualize models with *γ*. **H:** Association between fraction of iterations without updates to Δ_*k*_ and increase in *ν* for `fides ND/GN`. **I:** Association between change in average fraction of iterations where smallest eigenvalue *λ*_min_ of the transformed Hessian 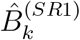 is negative and *γ* relative to `fides 2D/SR1` for `fides ND/SR1`

To better understand the performance differences between BFGS and SR1, we analysed the eigenvalue spectra of SR1 approximations and found that SR1 convergence rate *ν* was correlated with the average fraction of iterations where the transformed Hessian approximations 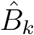 had negative eigenvalues (*r* = −0.71, *p* = 6.7 · 10^−3^, Fig 4E). This correlation suggests that directions of negative curvature approximated by the SR1 scheme tended not to yield good search directions; handling of negative curvature is therefore unlikely to explain observed improvements in convergence rates. In contrast to negative eigenvalues, we found that difference in *ν* between BFGS and SR1 was correlated with the average fraction of iterations in which the BFGS approximation did not produce an update (*r* = −0.88, *p* = 8.3 · 10^−5^, Fig 4F). The BFGS approximation is not updated when the curvature condition is violated (see Section 2.4). Such a high fraction of iterations not prompting updates is surprising, since the violation of the curvature condition is generally considered to be rare [13]. It is nonetheless a plausible explanation for lower convergence rates, since the BFGS approximation is not expected to always converge to the true Hessian under such conditions [13].

For three problems, the BFGS (orange arrows, Fig 4C) and/or SR1 (green arrows, Fig 4C) approximations increased *γ* (1.29 to 3.12 fold change; *Boehm*, *Crauste*, *Schwen*), but for most problems *γ* was reduced by more than two-fold (−∞ to 0.46 fold change; SR1+BFGS: *Beer*, *Fiedler Fujita*, *Isensee*, *Lucarelli Weber*, *Zheng*; BFGS: *Bachmann*; SR1: *Brannmark*), canceling the benefit of faster convergence rate for many of these problems. Paradoxically, we found that convergence count changes were correlated with changes in the average fraction of iterations having ill-conditioned trust-region subproblems (BFGS: *r* = 0.86, *p* = 1.6 · 10^−4^; SR1: *r* = 0.73, *p* = 4.8 · 10^−3^, Fig 4G). Therefore, improved conditioning of the trust-region subproblem, unexpectedly, came at the cost of smaller regions of attraction for minima having low objective function values.

We complemented the analysis of `2D` methods by evaluating their respective `ND` methods, which almost exclusively performed worse than `2D` methods. We found that `Fides ND/GN` outperformed `Fides 2D/GN` on two problems (1.43 to 3.25-fold change; *Crauste*, *Zheng*) and performed similarly on one problem (0.99-fold change; *Fiedler*). `Fides ND/BFGS` outperformed `Fides 2D/BFGS` on three problems (1.16 to 1.37-fold change; *Boehm*, *Fujita*, *Schwen*) and performed similarly on three examples (1.01 to 1.06-fold change; *Bachmann*, *Bruno*, *Fiedler*). `Fides ND/SR1` outperformed `Fides 2D/SR1` on two problems (1.51 to 1.70-fold change; *Crauste*, *Schwen*). These results were surprising, since the use of `2D` methods is generally motivated by lower computational costs, not better performance; the `ND` approach gives, in contrast to the `2D` approach, an exact solution to the trust-region subproblem. For GN, the change in convergence rate *ν* was correlated with the change in average fraction of iterations in which the trust-region radius Δ_*k*_ was not updated (*r* = −0.8, *p* = 1.1 · 10^−3^, Fig 4H). For `fides SR1/ND`, the change in *γ* with respect to `fides SR1/2D` was correlated with the change in average fraction of iterations in which 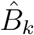 had negative eigenvalues (*r* = −0.72, *p* = 1.2 · 10^−2^, Fig 4I). This suggests that inaccuracies in Hessian approximations may have stronger impact on `ND` methods as compared to `2D` methods, thereby mitigating advantages that are theoretically possible.

Overall these results suggest that BFGS and SR1 approximations can improve optimization performance through faster convergence, but often suffer from poorer global convergence properties. Thus, they rarely outperform the GN approximation. BFGS and SR1 perform similarly on most problems, with the exception of a few problems for which BFGS cannot be updated due to violation of curvature conditions. We conclude that, while saddle points may be present in some problems, they do not seem to pose a major issues that can be resolved using the SR1 approximation.

### 3.4 Hybrid Switching Approximation Scheme

We hypothesized that the high success count *γ* of the GN approximation primarily arose in the initial phase of optimization, which determines the basin of attraction on which optimization will converge. In contrast, we surmised that the high convergence rate *ν* of the BFGS approximation was due more accurate Hessian approximation in later phases of optimization, when convergence to the true Hessian is achieved. To test this idea, we designed a hybrid switching approximation that initially uses a GN approximation, but simultaneously constructs an BFGS approximation. As soon as the quality of the GN approximation becomes limiting, as determined by a failure to update the trust-region radius for *n*_*hybrid*_ consecutive iterations, the hybrid approximation switches to the BFGS approximation for the remainder of the optimization run.

We compared the hybrid switching approach using different values of *n*_*hybrid*_ (25, 50, 75, 100) to `fides 2D/GN` (equivalent to *n*_*hybrid*_ = ∞) and `fides 2D/BFGS` (equivalent to *n*_*hybrid*_ = 0). Evaluating optimizer performance *ϕ*, we found that the hybrid approach was successful for all problems, with *n*_*hybrid*_ = 50 performing best, improving *ϕ* by an average of 1.51 fold across all models (range: 0.56 to 6.34-fold change). The hybrid approach performed better than `fides 2D/GN` and `fides 2D/BFGS` on 5 problems (1.71 to 6.34-fold change;*Crauste*, *Fiedler*, *Fujita*, *Lucarelli*, *Zheng*; + signs Fig 5A). It performed better than `fides 2D/GN`, but worse than `fides 2D/BFGS` only on one problem (*Boehm*; (+) sign Fig 5A). The hybrid approach performed similar to `fides 2D/GN` on 5 out of the 7 remaining problems (0.89 to 1.03-fold change; *Beer*, *Brannmark*, *Bruno*, *Isensee*, *Schwen*; = signs Fig 5A). Decomposing *ϕ* into *γ* and *ν*, we found that hybrid switching resulted in higher *ν* for the four problems in which `fides 2D/BFGS` had higher *ν* than `fides 2D/GN` (*Beer*, *Boehm*, *Fiedler*, *Zheng*), as well as three additional problems (*Crauste*, *Fujita*, *Lucarelli*). These were the same three problems for which SR1 had higher *ν* (Fig 4B), but BFGS did not, as a consequence of a high number of iterations without *B*_*k*_ updates (Fig 4F). Consistent with this interpretation, we confirmed that the hybrid approach had very few iterations without *B*_*k*_ updates. In contrast to BFGS, the hybrid switching approach maintained a similar *γ* as GN, meaning that higher *ν* generally translated into higher *ϕ*. Evaluating the overlap between start-points that yielded successful runs for GN showed a higher overlap for the hybrid switching approach as compared to BFGS (Fig 5D), suggesting that higher *γ* for the hybrid switching was indeed the result of higher similarity in regions of attraction. These findings further corroborate that local convergence of `fides 2D/GN` is slowed by the limited approximation quality of GN.

**Fig 5:**
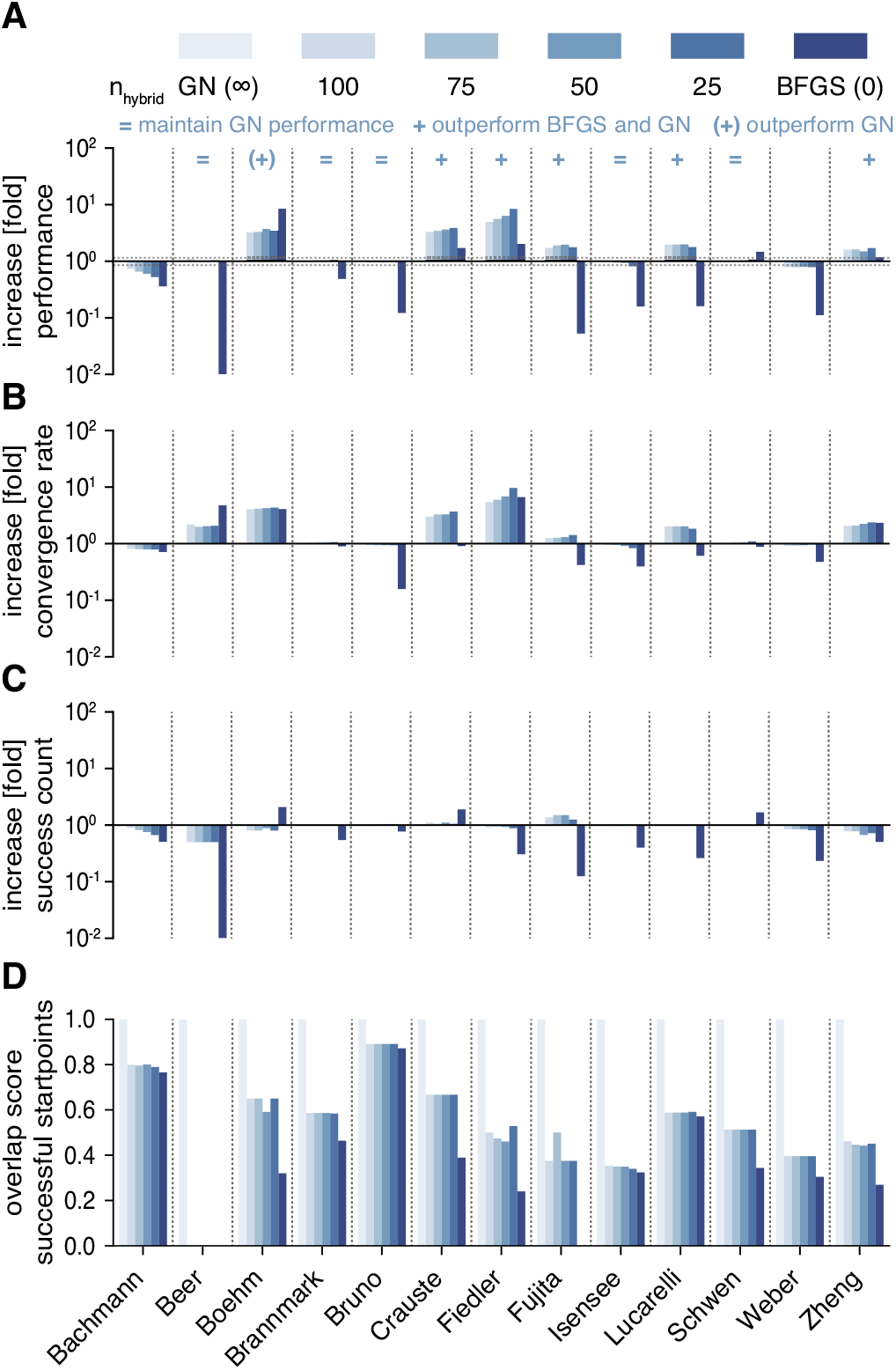
Evaluation of hybrid switching approximation. Color scheme is the same in all panels. All increases/decreases are relative to `fides 2D/GN`. **A:** Performance comparison (*ϕ*). **B:** Increase in convergence count *γ*. **C:** Increase in convergence rate *ν*. **D:** Overlap score for start-points that yield successful optimization runs with respect to `fides 2D/GN`.

**Fig 6:**
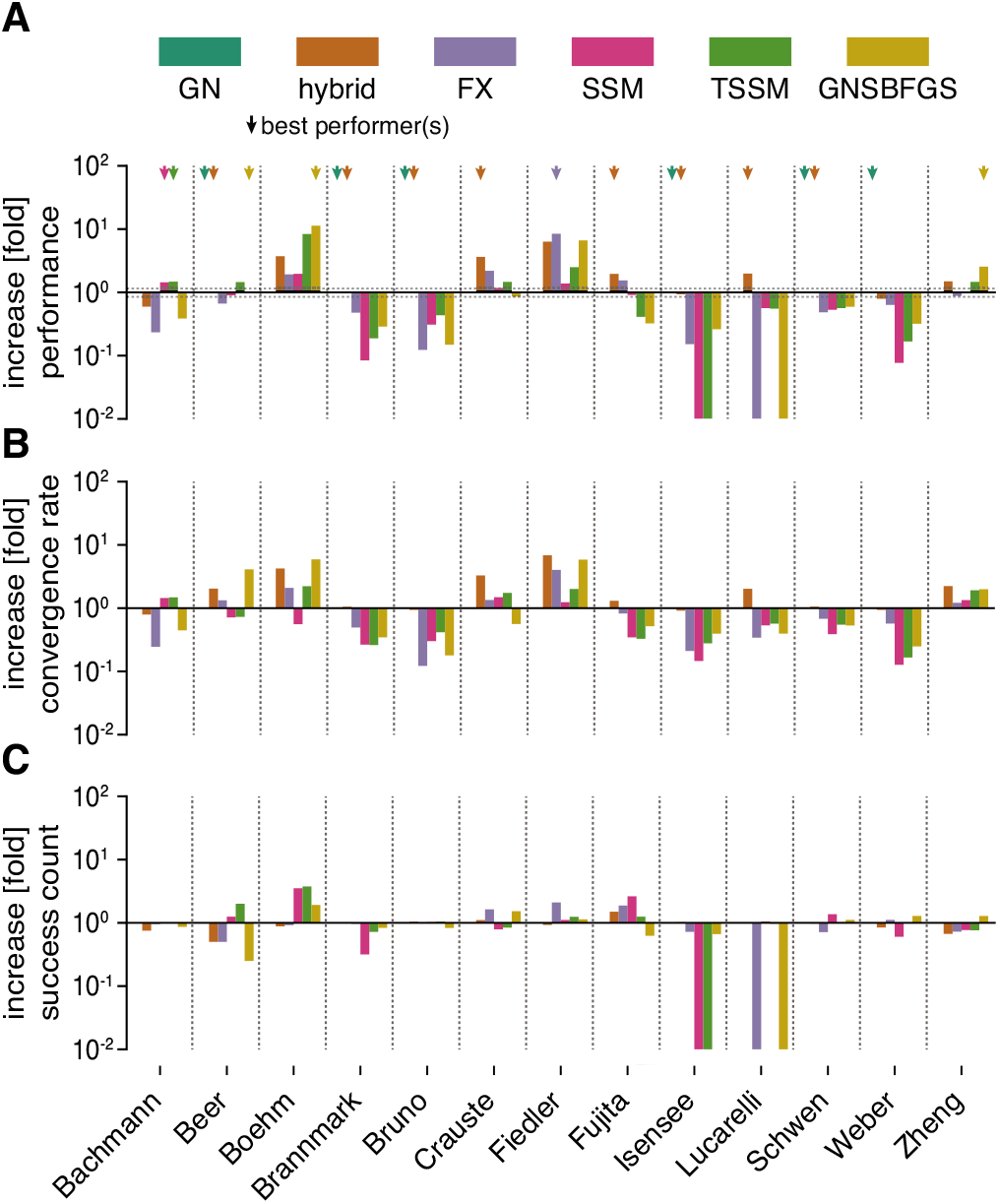
Evaluation of hybrid switching approximation. Color scheme is the same in all panels. All increases/decreases are relative to `fides 2D/GN`. **A:** Performance comparison (*ϕ*). **B:** Increase in convergence count *γ*. **C:** Increase in convergence rate *ν*.

## 4 Comparison of Hybrid Approximation Schemes

Inaccuracies in the GN approximation have been previously been discussed in the optimization literature and are known to lead to slow convergence and even divergence of optimization runs [33, 34]. Several methods have been proposed to address this issue in the context of non-zero residual problems. These include the Structured Secant Method (SSM) [36], the Totally Structured Secant Method (TSSM) [37], the hybrid scheme by Fletcher and Xu (FX) [39] and the Gauss-Newton Structured BFGS (GNSBFGS) approach [34]. All of these methods combine the GN and BFGS approximations in different ways (see Section 2.4).

We implemented support for these Hessian approximation schemes in `fides` and compared optimizer performance *ϕ* against `fides 2D/GN` and the best performing hybrid switching method (*n*_*hybrid*_ = 50). We found that the hybrid switching method was among the best performing methods (fold change-0.85 to 1.15) on a majority of problems (7 out of 13; *Beer*, *Brannmark*, *Bruno*, *Crauste*, *Fujita*, *Lucarelli*, *Schwen*) and was the only method other than `fides 2D/GN` that resulted in successful runs for all problems. `fides 2D/GN` was among the best performers on 6 problems (*Beer*, *Boehm*, *Bruno*, *Isensee*, *Schwen*, *Weber*) whereas GNSBFGS was among the best performers for three problems (*Beer*, *Boehm*, *Zheng*) and FX for one problem (*Fiedler*). Both GNSBFGS and FX failed for one problem (*Lucarelli*). SSM and TSSM were among the best performers on one problem (*Bachmann*) and failed on one problem (*Isensee*). We conclude that the hybrid switching method is the most reliable and efficient method among all methods that we tested.

## 5 Discussion

In this paper we evaluated trust-region methods implemented in `fides` against Python and MATLAB implementations of the same algorithm on 13 established benchmark optimization problems. Using the logging capabilities of `fides`, we were able to link success counts *γ* and convergence rate *ν* to numerical properties of optimization traces, spurring the development of a novel hybrid switching scheme that uses two approaches for Hessian approximation: the Gauss-Newton approximation early in a run (when the basin of attraction is being determined) and the BFGS approximation later in a run (when a fast convergence rate to the local minimum is crucial). For many problems, `fides` in combination with hybrid switching exhibited the best performance and resolved issues with inconsistent final objective function values. However, across all problems we studied, there was no single uniformly superior optimization method. Hybrid switching improved average performance and was superior on a majority of problems, but there remained a minority of problems where other hybrid methods performed substantially better. In general, this heterogeneity in optimizer performance is in line with the infamous “no free lunch” theorem of optimization [7], and suggests the existence of distinct problem classes that we so far were not able to identify. We anticipate that future innovation in optimization methods will likely be driven either by better understanding how performance differences relate to model structure, enabling *a priori* selection of best performing optimizers for specific problems, or to numerical properties of optimization traces, driving the development of new adaptive methods. Thus, the availability of multiple Hessian approximation schemes and trust-region subproblem solvers within the modular structure of `fides` will be of general utility both in optimizing specific models of interest and in future research on optimization methods.

Our findings suggest that previously encountered issues with `fmincon` and `lsqnonlin` are likely due to premature optimizer termination and not “rugged” objective function landscapes having many similar local minima. Our results corroborate previous findings from others showing that the use of Gauss-Newton approximations can be problematic for optimization problems featuring sloppy models. However, we did not find that methods designed to handle saddle points improved performance. The inconsistent and often poor performance of BFGS and SR1 approaches was unexpected, but our findings suggest that the problem arises in the global convergence properties of BFGS and SR1, as revealed by lower convergence counts *γ*. Global convergence properties depend on the shape of the objective function landscape and are therefore expected to be problem-specific. BFGS and SR1 may therefore perform better when combined with hybrid global-local methods such as scatter search [50], which substantially benefit from good local convergence [43], but are less dependent on global convergence. Moreover, SR1 and BFGS approaches enable the use of trust-region optimization for problems in wich the GN approximation is not applicable, such as when a non-Gaussian error model is used [27] or when gradients are computed using adjoint sensitivities [30].

Even though we re-implemented a previously described optimization algorithm, we observed pronounced differences in optimizer performance across benchmark problems. For many examples, observed differences in performance among theoretically identical algorithms were similar or larger in magnitude as differences observed with distinct variants of an algorithm. One reason for these differences may arise from how numerical edge cases are handled. For example, `fides` uses a Moore-Penrose pseudoinverse to compute the Newton search direction for the 2D subproblem solver, while `fmincon` uses damped Cholesky decomposition and warns the user in cases of ill-conditioning. Another possible source of difference is the use of different simulation and sensitivity computation routines. While both data2dynamics and AMICI employ CVODES [26] for simulation and computation of parameter sensitivity, there may be slight differences in implementation of advanced features such as handling of events and pre-equilibration. Overall, these findings demonstrate the complexity of implementing trust-region methods and the impact of subtle differences in numerical methods on optimizer performance. Thus, the benchmarking of algorithms requires consistent implementations.

Overall, our results demonstrate that `fides` not only finds better solutions to parameter estimation problems when state-of-the-art algorithms fail, but also performs on par or better on problems where established methods find good solutions. We expect that the strong performance shown in our work will generalize to other optimization problems beyond ODE models. Thus, we expect that the modular and flexible implementation of `fides` will drive widespread adoption within and outside the field of systems biology.

## Acknowledgements

This work was supported by the Human Frontier Science Program (Grant no. LT000259/2019-L1; F.F.), and the National Cancer Institute (Grant no. U54-CA225088; P.K.S). We thank the O2 High Performance Compute Cluster at Harvard Medical School for computing support and Carolin Loos, Daniel Weindl, Dilan Pathirana, Elba Raimundez, Erika Dudkin, Jan Hasenauer and Leonard Schmiester for providing Benchmark examples in PEtab format; we also thank Jan Hasenaur and Edward Novikov for feedback on the manuscript and Daniel Weindl for feedback on the implementation of `fides`.

